# Quantitative assaying of SpCas9-NG with fluorescent reporters

**DOI:** 10.1101/2022.08.04.502727

**Authors:** Alexandre Baccouche, Kevin Montagne, Nozomu Yachie, Teruo Fujii, Anthony Genot

**Affiliations:** Laboratory for Integrated Micro and Mechatronic Systems, CNRS-IIS UMI 2820, The University of Tokyo, Tokyo, 153-8505, Japan; Department of Mechanical Engineering, School of Engineering, The University of Tokyo, Tokyo, 113-8656, Japan; School of Biomedical Engineering, Faculty of Applied Science and Faculty of Medicine, The University of British Columbia, Vancouver, British Columbia, BC V6T 1Z3 Canada

## Abstract

The Cas9 enzyme has revolutionized biology in less than a decade. Engineering Cas9 to expand its functionality has become a major research goal, yet assaying variants of Cas9 remains a laborious task that is commonly performed with gel electrophoresis. Fluorescence assays have been reported for Cas9 but their utility for assaying variants of Cas9 has not been investigated in detail. Here we use a simple fluorescent assay to resolve differences of activity between the wild type Streptococcus pyogenes Cas9 (SpCas9) and SpCas9-NG, a variant with an expanded PAM repertoire. We compare the kinetics of the two enzymes on dozens of mutated RNA guides – highlighting the benefits of fluorescence such as quantitativity, sensitivity, multiplexing, non-invasiveness and real-timeness. This validates fluorescence as a tool for engineering Cas9 and lays the groundwork for directly evolving Cas9 in microfluidic compartments.

## INTRODUCTION

In less than a decade, the CRISPR-Cas9 system^1,2^ has upended many fields of life sciences. Its programmability has enabled genome editing, synthetic gene regulation, or live imaging of chromosomes^3–6^, and it augurs groundbreaking advances for human healthcare^7^, agricultural biotechnology^8^ or molecular diagnosis^9^.

The CRISPR-Cas9 system comprises a Cas9 enzyme and a guide RNA (gRNA), itself composed of a conserved region (the scaffold, which initiates binding with Cas9), and a variable region (the spacer, which directs Cas9 to the target region for cleavage). Upon loading the guide RNA, the Cas9:gRNA complex first scouts double-stranded DNA (dsDNA) for a specific sequence (the Proto Adjacent Motif or PAM). Once it finds one, it unwinds the DNA nearby and attempts to match it with the spacer of the guide RNA by forming an R-loop. When the two domains match, Cas9 cleaves both sides of the target DNA (3 base pairs upstream of the PAM), leaving a double-stranded break in the genome that is the entry point for genome editing.

Protein engineers have expanded the functionality of Cas9^10–14^. The simplest and most common mutation consists in deactivating one of the endonuclease lobes (producing a programmable nicking enzyme) or both lobes (producing the so-called dead Cas9, which has lost its ability to cut DNA but retains its ability to lock onto target DNA). Splitting Cas9 is also a common strategy to control the activity of Cas9^15,16^. The enzyme is split in two nonfunctional parts that become functional when reunited by a chemical or physical signal (small molecule, light, temperature..).

A highly sought-after variant is a Cas9 with an expanded PAM repertoire. Wild type (WT) Cas9 recognizes a 3-letter motif NGG, which means that a GG sequence must be located 1 bp upstream of the domain recognized by the spacer. This constraint on the PAM limits the repertoire of targettable sequences, as for instance GG/CC motifs comprise only ~ 5% of the human genome. Thanks to rational protein engineering, Cas9 enzymes with an expanded NG PAM have been reported ^7–9^. In the SpCas9-NG variant, the base-specific interactions between Cas9 and the third G of the PAM were deleted, and the surrounding amino residues were tuned to compensate for the loss of stability. These NG variants considerably expand the repertoire of targetable sequences, since G/C nucleotides comprise about ~41% of the human genome^17^.

Assaying the activity of Cas9 is a bottleneck in the cycle of protein engineering. It is commonly done by incubating together the Cas9 variant, a guide RNA, and a target plasmid, and then resolving the cleaved plasmid by gel electrophoresis^1^. But this method is laborious, poorly adapted to the ubiquitous 96-well format (a gel slab accommodates at most a dozen wells) and prone to failures (smeared bands, buffer leakage, gel tearing or uneven staining occur commonly in the hands of unskilled operators). Gel electrophoresis also lacks multiplexing: one separate tube and gel lane must be allocated for each PAM to be tested. Additionally gel electrophoresis is an end-point method that lacks temporal resolution, sensitivity and quantitativity. Lastly the readout of electrophoresis is incompatible with modern tools for protein evolution, like the droplet-based directed evolution of protein with microfluidics, which use fluorescence in their screening step^18^. These drawbacks make gel electrophoresis poorly adapted to high-throughput and automated screening of variants of Cas9.

Fluorescence assays have many benefits that gel electrophoresis lacks. Fluorescence is real-time, multiplexed, sensitive and quantitative. An unskilled operator can easily collect hundreds of fluorescent traces in a day with automated instruments (plate reader, spectrophotometer, thermocyclers...), standardised plasticware and pipettes (e.g. 96-well or 384-well plates), as well as commercial services for synthesis of fluorescent DNA probes (hundreds of commercial dyes are now available). This throughput is crucial for quantitative fields like enzymology (which tests dozens or hundreds of combinations of experimental conditions like concentration, salinity, temperature...), or molecular diagnostics (which requires testing hundreds of patient samples per day). Fluorescence has revolutionised many areas of life science - replacing venerable but hazardous methods like radioactive quantification. Real-time measurement has also enabled PCR in its modern and quantitative form (qPCR). At the forefront of development, new methods like DNA paint take advantage of fluorescently labelled nucleic acids to push forward super-resolution microscopy ^19^.

In the past years, a few groups have introduced fluorescent assays for Cas9. Mekler and colleagues reported a simple yet robust displacement-based assay to follow in real-time the binding of Cas9 to its target and the formation of the R-loop ^20–22^. They used this assay to evaluate the competitive binding of nonspecific RNA to Cas9 - which was hypothesised to be one reason why Cas9 usually shows decreased activity when operated *in vivo*. This fluorescent assay was used to prototype allosteric RNA guides ^23,24^. Two other reports used high-throughput fluorescent assays to sift through massive chemical libraries with hundreds of thousands of compounds and identify small molecule inhibitors of Cas9^25,26^. Lastly, indirect fluorescent assays have also been reported - though they do not directly yield the kinetics of Cas9 ^27,28^.

Fluorescence assays could speed up the engineering or evolution of Cas9 proteins. The throughput of fluorescence could massively accelerate the assaying of variants, while the quantitativity of fluorescence could resolve incremental changes in activity, which would otherwise be lost with semi-quantitative methods like gel electrophoresis. Lastly, fluorescence is more sensitive than conventional gel electrophoresis, which could reduce the amount of enzyme needed down to scales that are easily produced with cell-free protein expression^27^. Yet reports on the utility of fluorescence for assaying variants of Cas9 have been elusive. Here we use a fluorescence assay to systematically compare the activity of the SpCas9-NG variant against the wild type enzyme. We compare the response of the two enzymes on four PAMs, and with RNA guides of varying lengths and secondary structures. Lastly, we multiplex the assaying of Cas9 - comparing in one pot the activity of the Cas9 enzymes on four PAMs. Overall, our results show that fluorescence assays are a powerful tool to assay the activity of Cas9 mutants, and their dependence on guide sequence.

## MATERIALS AND METHODS

### Reagents

SYBR Green II RNA gel stain was purchased from Thermo Fisher Scientific (Eugene, OR, USA). Propan-2-ol and dithiothreitol were from Sigma-Aldrich (Saint-Louis, MO, USA). Tris-EDTA buffer (TE) and glycerol were from Nacalai Tesque (Kyoto, Japan). Ribonucleotide triphosphates (rNTP, N0450), T7 RNA polymerase (M0251) and murine RNase inhibitor (M0314) were obtained from New England Biolabs (NEB, Ipswich, MA, USA). gBlock Gene Fragments were purchased from Integrated DNA Technologies (IDT, Coralville, IA, USA). Other oligonucleotides were HPLC-purified and obtained from Biomers (Ulm, Germany). All sequences are listed in Supplementary Table 1. To enable specific monitoring of beacon cleavage by Cas9, the bottom strands of the beacons were conjugated at their 3’ end to the fluorophores Atto 532, Atto 590, Atto 647N or DY-681, whose maximum absorption/emission wavelengths are respectively 532/554, 594/624, 646/664, and 691/708 nm. The top strands were conjugated at their 5’ end to various quenchers: BMN-Q535 for Atto 532, BMN-Q620 for Atto 590, BBQ-Q650 for Atto 647N and BMN-Q620, BMN-Q650 or BBQ-650 for DY-681. All nucleotides were dissolved in TE.

### Cas9 synthesis and purification

The Cas9 proteins were expressed as previously reported ^11^.

### Guide RNA transcription

RNA guides were transcribed from gBlock Gene Fragments. Transcription was carried out in a 25 μL volume containing 40 mM Tris-HCl, pH 7.9, 6 mM MgCl_2_, 1 mM DTT, 2 mM spermidine, 5000 units.mL^-1^ T7 RNA polymerase, 500 μM of each rNTP, 1000 units.mL^-1^ murine RNase inhibitor, SYBR Green II 10 nM (1x), and 4 nM gBlock. The reaction was run for 7 h at 34 °C on a CFX96 real-time PCR detection system (Biorad) and monitored thanks to the SYBR Green II signal.

### Guide RNA purification

Transcribed RNA was purified by adding 280 μL of nuclease-free water to the 20 μL transcription reaction, followed by 30 μL of 3 M sodium acetate, mixing, and adding 330 μL of propan-2-ol, thus precipitating the RNA. The RNA was then pelleted by centrifugation at 10.000 g for 10 min at 4°C. The RNA pellet was washed once in 75% ethanol, centrifuged at 10.000 g for 5 min at 4°C, air dried and resuspended in nuclease-free water before quantification with a Biodrop spectrophotometer (Berthold Technologies).

### Reaction assembly

Reactions were assembled in a buffer containing 20 mM HEPES, pH 7.5, 100 mM KCl, 2 mM MgCl_2_, 1 mM DTT, 5% glycerol, and 1000 units.mL^-1^ murine RNase inhibitor. All reagents except the double-stranded beacon were first assembled in a total volume of 9 μL and run at 37 °C in a CFX96 real-time PCR detection system (Biorad). Both WT SpCas9 and SpCas9-NG are highly active at this temperature; the activity is relatively stable between 30 and 45°C but drops suddenly above 46°C for SpCas9-NG and above 48°C for WT SpCas9 (Supplementary Figure 4). After about 5 minutes to allow stabilization of the temperature, 1 μL of beacon solution was injected in each tube, giving a final volume of 10 μL, while the machine was paused for a minimal period. Unless specified otherwise, the final concentration of beacon was 40 nM, that of Cas9 (WT or NG) was 200 nM, and that of guide RNA was 400 nM. After cleavage of the beacon, the cleaved strands separate thus releasing the fluorophore on the bottom strand from the vicinity of the quencher. Cleavage of the beacon was thus monitored specifically by measuring the fluorescence signal of the beacon’s bottom strand fluorophore (Fig.1 top). Fluorescence signals were acquired every 15 s by the CFX96 real-time PCR detection system in the channels corresponding to the fluorophore of the beacon’s bottom strand.

**Figure 1.**
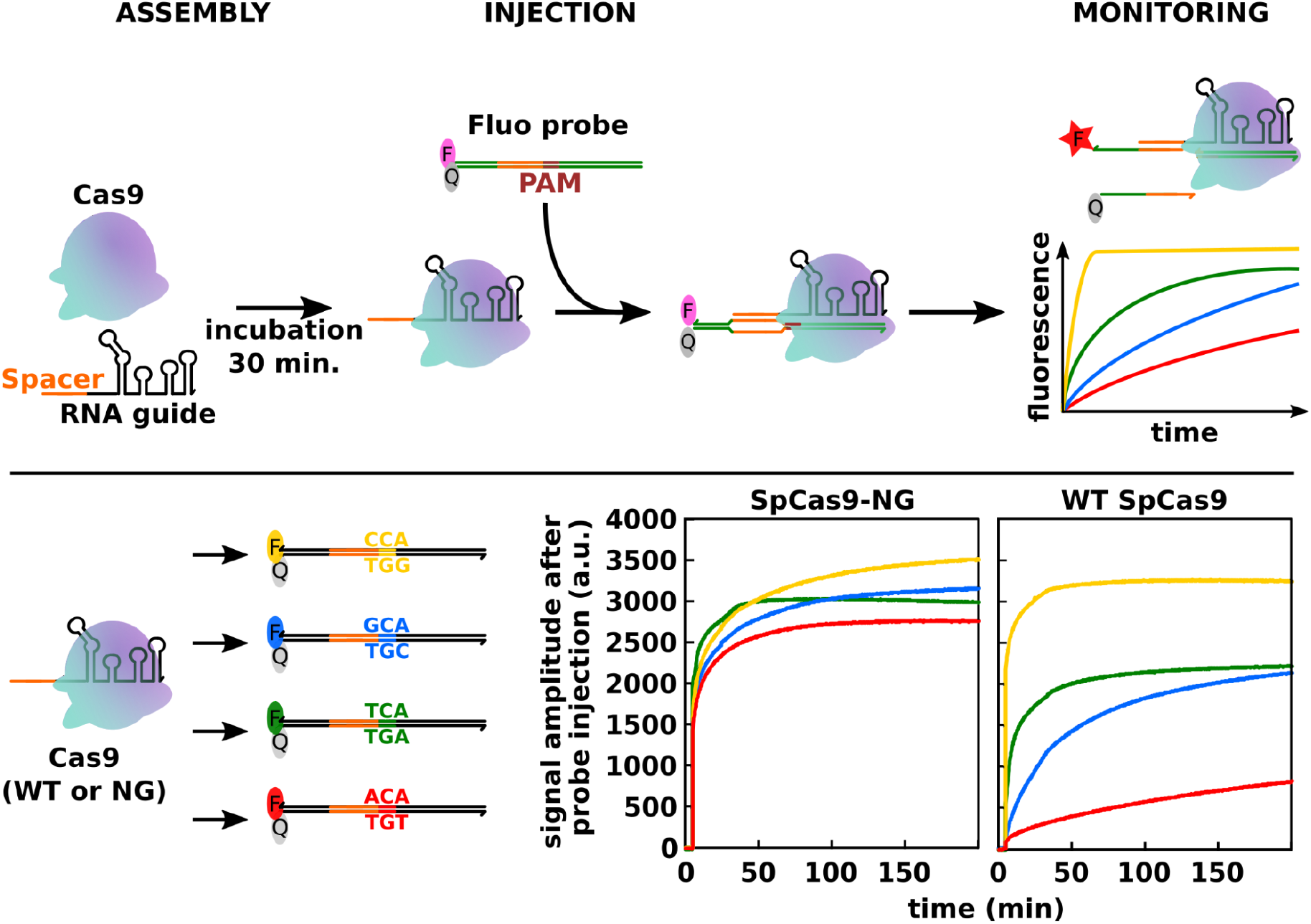
Fluorescence assaying of Cas9. (Top) Workflow of the cleavage assay. The reporter is a dsDNA strand bearing a pair of fluorophore and quencher, and which is targeted by the spacer of an RNA guide. After pre-incubation of the guide and Cas9, this complex is mixed with the reporter. After recognizing the PAM, Cas9 unwinds the adjacent domain and matches it with the spacer. This R-loop separates the fluorophore from its quencher, and thus increases the fluorescence. (Bottom) Assaying of SpCas9-NG or WT SpCas9 (right) against four distinct PAMs.

## RESULTS

We use the fluorescent reporter of Mekler et al. to assay the activity of Cas9 and its variants ^20^. Briefly, the reporter is made of a DNA duplex (called beacon here) with a PAM sequence and two moieties at one of its ends (a fluorophore and a quencher). A Cas9 loaded with a guide recognizes the PAM sequence of the reporter and initiates an R-loop - attempting to match the sequence of the spacer with that of the reporter. This invasion by the spacer separates the two ends of the duplex, increasing fluorescence. Of note, this reporter does not directly report on the cleavage of the dsDNA beacon by Cas9, but on its invasion by the RNA spacer (as demonstrated by the fact that it also reports activity with dCas9).

We first verified that the fluorescent assay corroborates the cleavage kinetics of WT SpCas9 and SpCas9-NG as measured by gel electrophoresis (Fig. 1 of Nishimasu et al.^11^). Both gels and fluorescence measurements rank the kinetics of WT SpCas9 on PAM as TGG>>TGA>TGC>TGT. Gels and fluorescence measurements also agree that SpCas9-NG is active on all presented PAMs, and that SpCas9-NG is slightly slower than WT SpCas9 on the canonical PAM TGG. This agreement between gel and fluorescence kinetics suggests that finding a PAM and forming an R-loop are the rate limiting steps, rather than the cleavage of DNA (which is not directly measured by the fluorescent reporter ^20^.

We used the fluorescent assay and the thermal gradient of a thermocycler to quickly evaluate the temperature stability of the Cas9 enzymes (Supplementary Fig. 4). Both enzymes are stable up to ~45°C, at which point their activities drop precipitously with increasing temperatures (although WT SpCas9 was marginally more stable than SpCas9-NG over 45°C). This thermal stability is in line with a previous study with differential scanning calorimetry which found that WT SpCas9 denatured around 45°C ^29^.

The direct delivery to cells of RNA guides complexed with Cas9 is one of the main modalities for editing genes. In order to improve delivery, there is interest in minimizing the size of this cargo, either with more compact CRISPR nucleases^30–32^, or by removing superfluous domains in the guide^33^. It is essential to know how much of the guide can be truncated while retaining its function. Truncation in the variable domain of the guide (the spacer) has been extensively studied - mainly for the purpose of reducing off-target effect^33^. Yet, truncation in the constant domain of the guide (the scaffold, i.e the region located downstream of the spacer) has been comparatively less studied.

We used the fluorescent assay to test guides of various lengths, progressively truncating the guide from its 3’ end (Fig. 2). SpCas9-NG and WT SpCas9 show a similar, yet slightly distinct, dependence on the length of the guide. The activity of both Cas9 enzymes is suppressed for guides shorter than 62 bases, *i.e*. both enzymes require the presence of the stem and the nexus - a small but crucial secondary structure that contacts the α-helical and nuclease lobes in Cas9 ^15^. In WT SpCas9, those two structural domains are the minimal requirement for activity^34^, and guides longer than 66 bases show strong activity with WT SpCas9 (Fig. 2). However SpCas9-NG needs an extra domain, the first 3’ hairpin (located in the 70-80 nt range) to show an appreciable activity (Fig. 2). Once these 3 domains (stem, nexus and first 3’ hairpin) are present in the guide, SpCas9-NG is strongly active on all 4 PAMs - while WT SpCas9 is only fully active on the TGG PAM, and to a lesser extent on the TGA PAM. Similar results were obtained using another beacon with a different spacer sequence (Supplementary Figure 1).

**Figure 2.**
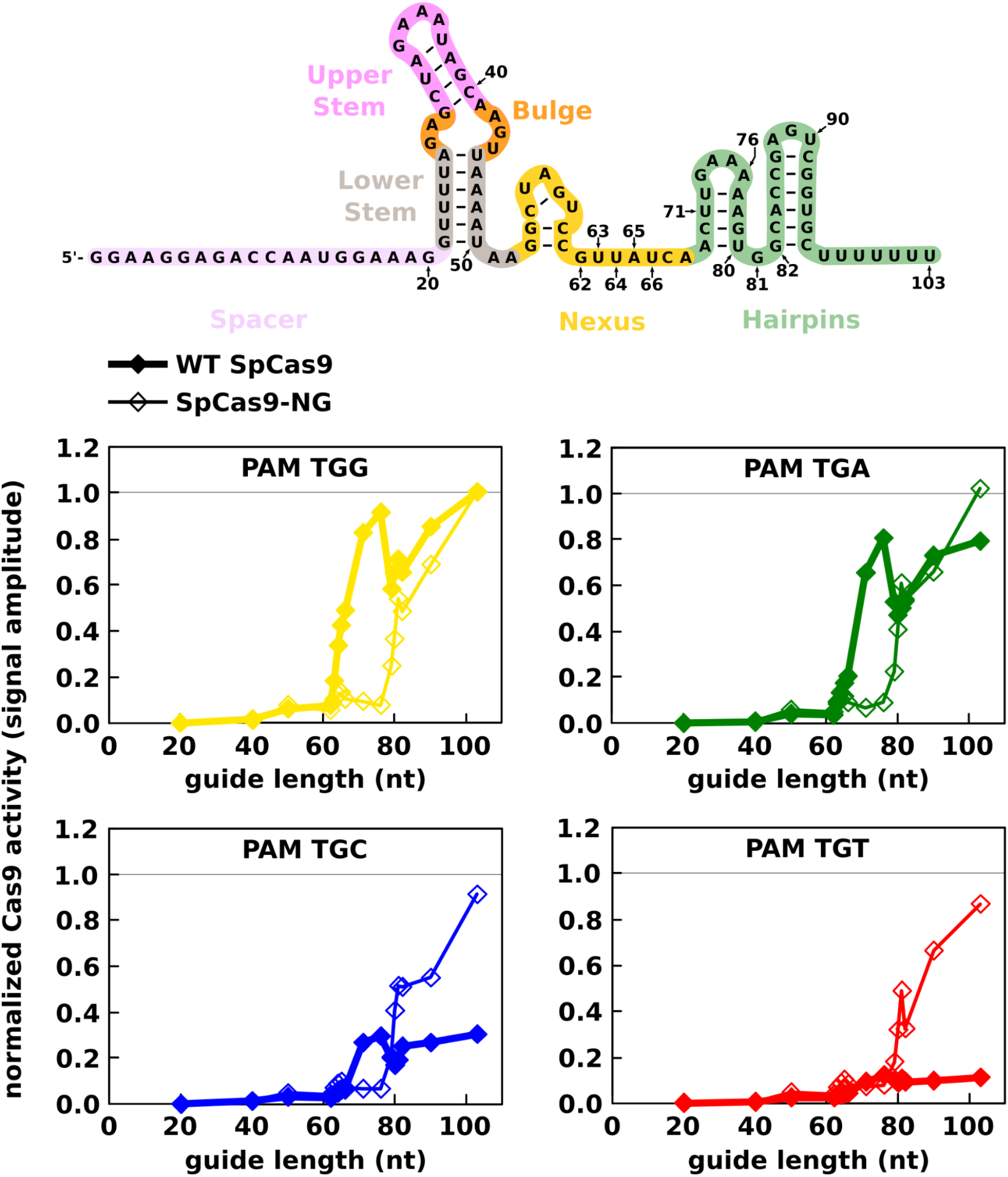
Effect of the guide’s length on the Cas9 activity. (top) anatomy of the guide RNA showing the spacer, upper and lower stem, bulge, nexus and hairpins with the RNA sequence. (bottom) WT SpCas9 (thick line, filled diamond) or SpCas9-NG (thin line, open diamond) was first incubated with guide RNAs of different lengths. The top schematic diagram shows the full-length guide RNA sequence; the shorter guide RNAs all contain the spacer (in mauve) and finish at one of the positions indicated in the diagram. After stabilization of the fluorescence signal, the beacon (beacon 3 containing one of four PAMs) was then injected and the signal amplitude was measured 10 min after injection. The signal amplitudes were normalized, for each Cas9, to the value obtained with the full-length 103 bp-long guide RNA and the canonical beacon (with the PAM TGG).

We then assayed how the Cas9 enzymes responded to mutations in the guide. There is an increasing interest in engineering the secondary structure of the guide to control the activity of Cas9. For instance, the decoration of the upper stem with an aptamer helps to recruit transcription factors - converting Cas9 into a synthetic transcription factor^35^. DNA nanotechnologists have also explored the allosteric control of the guide with strand displacement - which converts the guide between active and inactive secondary structures^24,36,37^. Thus it is important to know which mutations in the guide are tolerated by Cas9 enzymes in order to expand the design space.

To that end, we selected a range of representative mutations that have been tested *in vitro* and *in vivo*^34^. We mutated the lower stem, the bulge, the nexus (Fig.3) or the 3’ hairpins (Fig. 4) - and measured the activity of SpCas9-NG and WT SpCas9, ten minutes after beacon injection. Overall, the assay confirms that SpCas9-NG and WT SpCas9 tolerate and reject the same kind of mutations in the guide. Mutations that conserve the secondary structure of the guide are tolerated to a certain degree (*e.g*. guide v3 coding for a mutation in the stem, and guide v15 coding for a mutation in the nexus). Mutations or deletions that disrupt crucial secondary structures like the bulge (guide v12) or the nexus (guide v18) completely abolished the activity of both enzymes. Similar results were obtained on a beacon with a different sequence (Supplementary Figure 3) or with shorter guides with mutations in the stem or nexus (though Cas9 seemed more sensitive to stem mutations with shorter guides, Supplementary Figure 2). On the other hand, deletions in the terminal 3’ hairpins, or even deletion of the first hairpin, had little effect on the activity of WT SpCas9 or SpCas9-NG (Fig. 4). Interestingly, mutations are symmetric with respect to the PAM: all PAMs are similarly affected by mutations, which confirms that the contacts of Cas9 with the guide are essentially uncoupled from the contacts of Cas9 with the PAM^11^.

**Figure 3.**
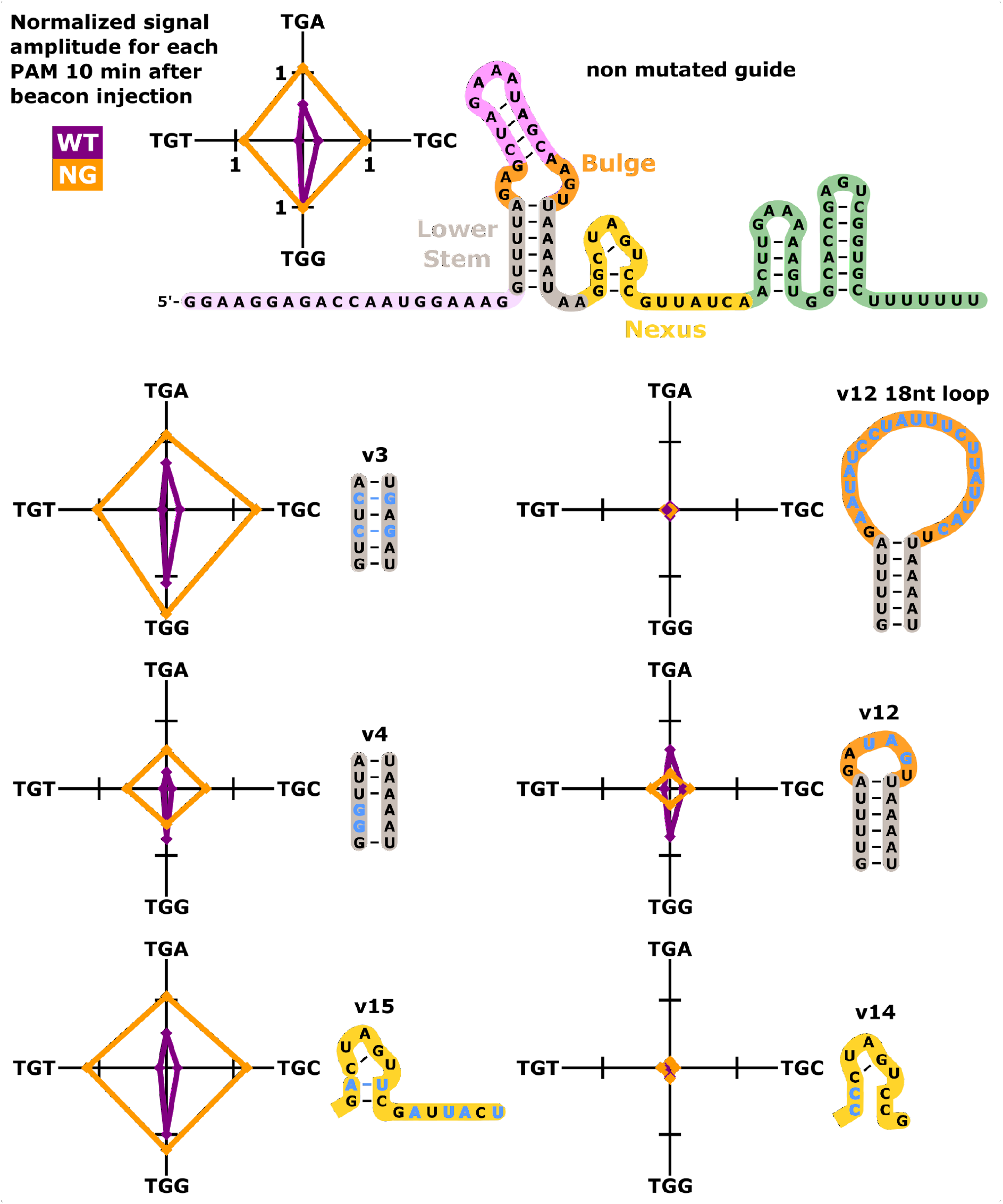
Effect of mutations in the guide RNA’s stem or nexus on the Cas9 activity. WT SpCas9 or SpCas9-NG was first incubated with full-length non-mutated guide RNA (top) or guide RNAs containing various mutations shown in blue in the schematic diagrams. The beacon (beacon 3 containing one of four PAMs) was then injected and the signal amplitude was measured 10 min after injection. The signal amplitudes were normalized, for each Cas9, to the value obtained with the full-length non-mutated guide RNA and the canonical beacon (with the PAM TGG). The radar charts show the normalized activity (signal amplitude) of WT SpCas9 (purple) or SpCas9-NG (orange) in the presence of each guide RNA depending on the PAM of the injected beacon. The nomenclature follows Briner et al.^34^

**Figure 4.**
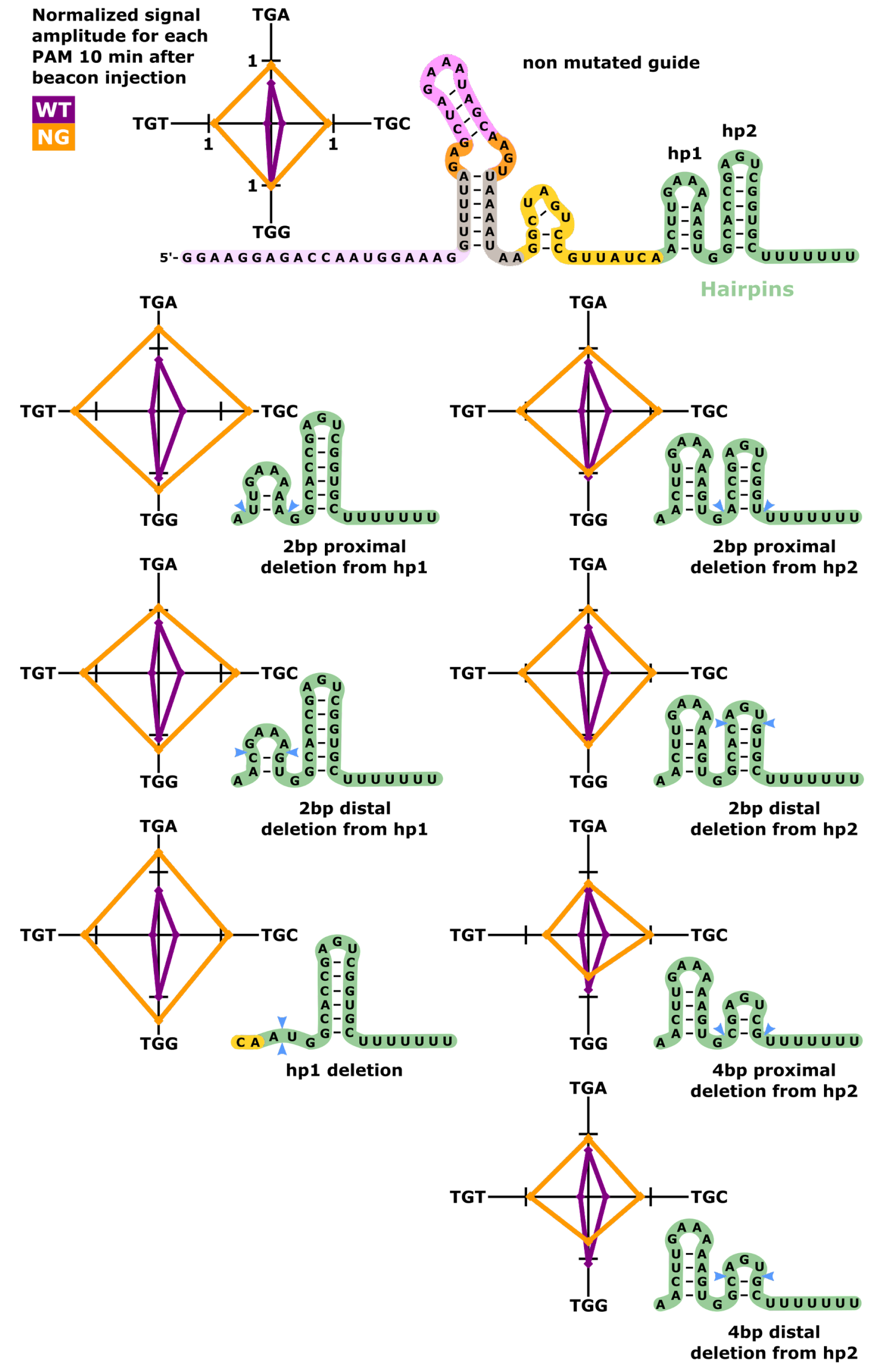
Effect of mutations in the guide RNA’s 3’ hairpins on the Cas9 activity. WT SpCas9 or SpCas9-NG was first incubated with full-length non-mutated guide RNA (top) or guide RNAs containing various deletions in the 3’ hairpins (hp1 or hp2) as shown in the schematic diagrams (the deletion sites are indicated with blue arrows). After stabilization of the fluorescence signal, the beacon (beacon 3 containing one of four PAMs) was injected and the signal amplitude was measured 10 min after injection. The signal amplitudes were normalized, for each Cas9, to the value obtained with the full-length non-mutated guide RNA and the canonical beacon (with the PAM TGG). The radar charts show the normalized activity (signal amplitude) of WT SpCas9 (purple) or SpCas9-NG (orange) in the presence of each guide RNA depending on the PAM of the injected beacon.

We then showcased the benefits of fluorescence for multiplexing measurements and speeding up workflows in protein engineering. We simultaneously measured the activity of Cas9 enzymes on four competing beacons (Fig .5). Each beacon bears a distinct PAM and reports its invasion by Cas9 in its own fluorescence channel. Multiplex measurements confirm the trends observed with singlex measurements (i.e. on single PAM). SpCas9-NG is equally active on all PAMs, as shown by the even distribution of activity in the four different fluorescence channels (right bars of Fig. 5). And as expected, WT SpCas9 is mostly active on the TGG PAM, and to a lesser extent on the TGA PAM (left bars). The pattern also suggests some non-linear competition between the PAMs for WT SpCas9. Overall, these results confirm that multiplex fluorescence measurements can be performed on Cas9, although no more than 4-6 targets can in principle be followed by fluorescence due to spectral overlap.

**Figure 5.**
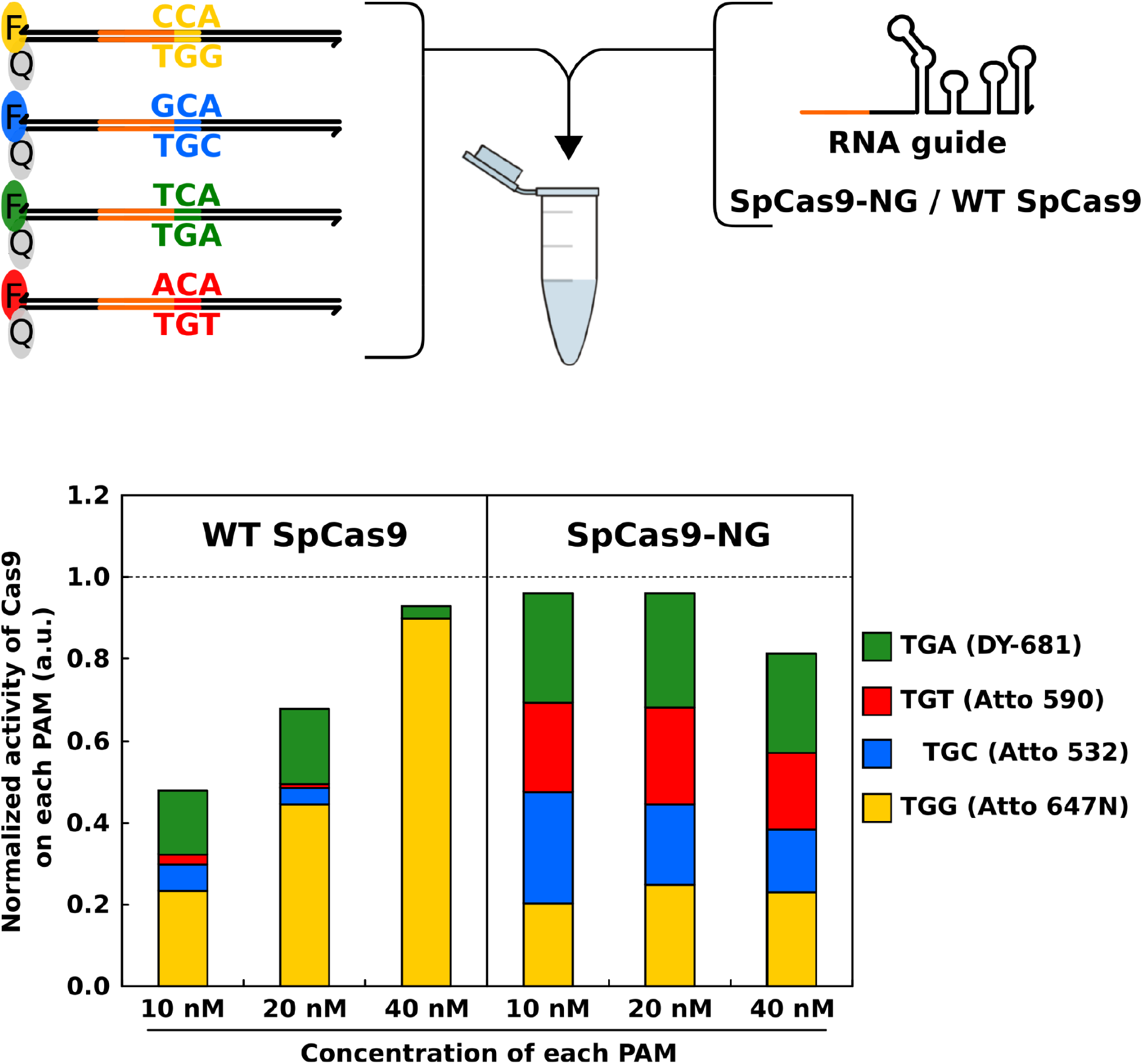
Multiplexed assessment of Cas9 activity on different PAMs. WT SpCas9 or SpCas9-NG was first incubated with full-length guide RNA. Then, four beacons - each with a different PAM and conjugated to a different fluorophore - were injected simultaneously. Cas9 activity was assessed by measuring the signal shift from each fluorophore 10 min after injection. The signal amplitudes were normalized to the value obtained after injecting on its own 40 nM of beacon 2 with the respective fluorophore (singleplex measurement). The stacked bar graph shows the activity of WT SpCas9 or SpCas9-NG on the different beacons present in solution at different concentrations.

## DISCUSSION

We have explored fluorescence as a modality for assaying an engineered variant of Cas9. Fluorescence reporting not only corroborates conventional gel electrophoresis, but it also gives a crispier picture of enzymatic kinetics. The sensitivity and quantitativity of fluorescence help to capture fine variations in activity which would otherwise be lost with gel electrophoresis. The resolution of incremental changes of activity should enable finer cycles in protein engineering, and help to home in on mutations that drive those changes in activity. Fluorescence measurements also require significantly less labour or starting materials than gel electrophoresis - providing a turnkey method to quickly screen Cas9 variants or more generally other types of CRISPR enzymes (Cas12, Cas13....).

Fluorescent assays could also be essential for the directed evolution of Cas9 in droplet microfluidics, whose massive parallelism and low consumption of reagents make it a powerful tool for protein engineers ^38–41^.

## DATA AVAILABILITY

The data that support the findings of this study are available on request from the corresponding author..

## ACKNOWLEDGEMENT

We warmly thank Pr Nishimasu and Pr Nureki (University of Tokyo) for kindly providing us with sample of wild type Cas9 and Cas9-NG. This work was supported by the JSPS Core-to-Core Program A (grant number:JPJSCCA20190006)

## FUNDING

This project is funded by the Japanese Society for Promotion of Science [JSPS KAKENHI JP17F17796], the French National Research Agency [ANR-17-CE18-0013], and by the Japanese Ministry of Education, Culture, Sports, Science and Technology through the Global Leader Program for Social Design and Management.

## CONFLICT OF INTEREST

AJG, TF and AB have filed a patent on Smart Guide RNA WO 2021/111641 A1.

