## Supplementary Material for "Quantitative assaying of SpCas9-NG with fluorescent reporters"

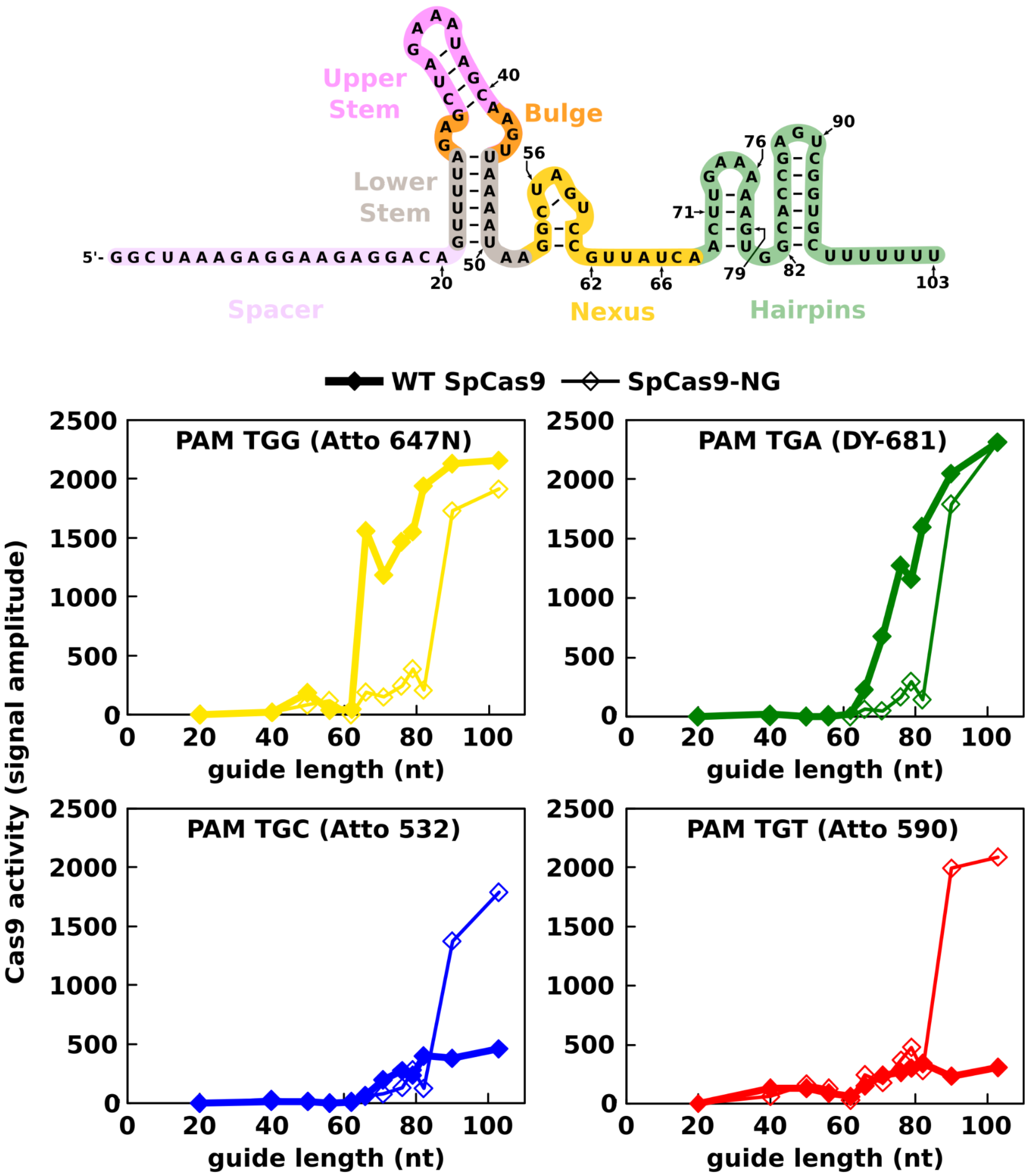

Supplementary Figure 1. Effect of the guide's length on the activity of Cas9 in the presence of beacon 2. WT SpCas9 (thick line, filled diamond) or SpCas9-NG (thin line, open diamond) was first incubated with guide RNAs of different lengths. The top schematic diagram shows the full-length guide RNA sequence; the shorter guide RNAs all contain the spacer (in mauve) and finish at one of the positions indicated in the diagram. After stabilization of the fluorescence signal, the beacon (beacon 2 containing one of four PAMs) was then injected and the signal amplitude was measured 10 min after injection. The plots show the signal amplitudes obtained depending on the beacon's PAM.

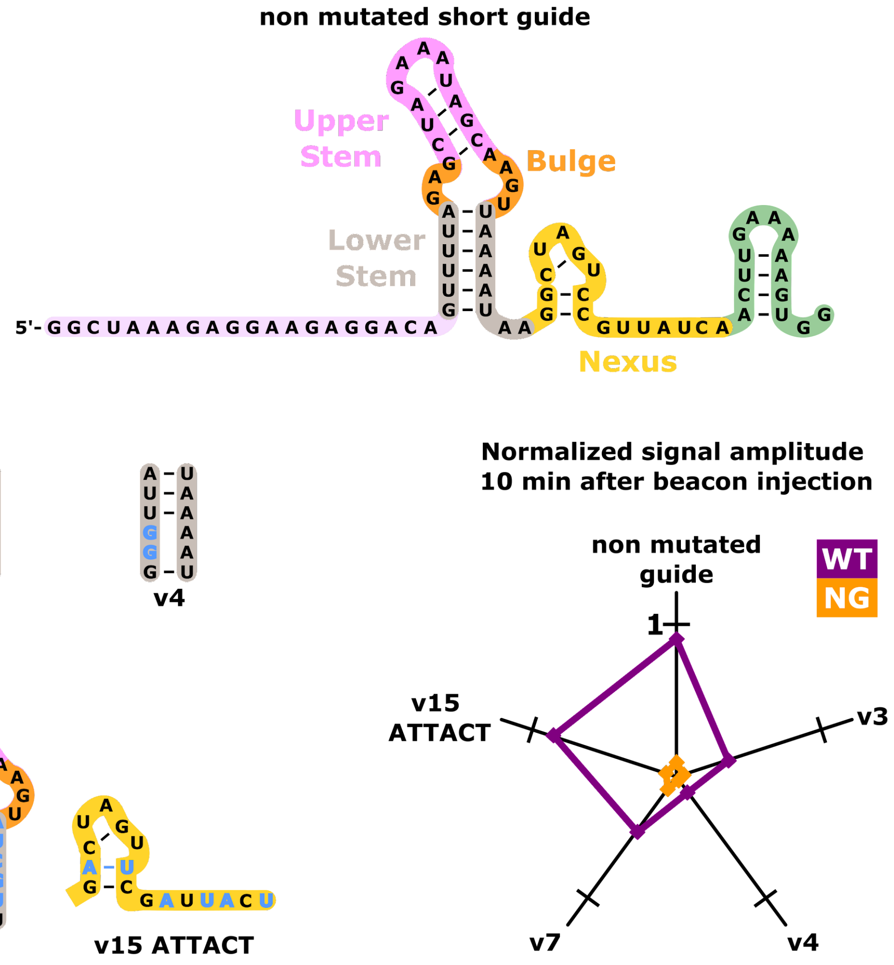

Supplementary Figure 2. Effect of mutations in the guide RNA's stem or nexus on the Cas9 activity. WT SpCas9 or SpCas9-NG was first incubated with a short (82 bp in length) non-mutated guide RNA (top) or guide RNAs containing various mutations shown in blue in the schematic diagrams (the naming refers to the convention of Barrangou et al.). After stabilization of the fluorescence signal, the canonical beacon 2 (containing the PAM TGG) was injected and the signal amplitude was measured 10 min after injection. The signal amplitudes were normalized to the value obtained with WT SpCas9 in the presence of full-length 103 bp-long non-mutated guide RNA and the canonical beacon 2. The radar chart shows the normalized activity (signal amplitude) of WT SpCas9 (purple) or SpCas9-NG (orange) in the presence of each guide RNA.

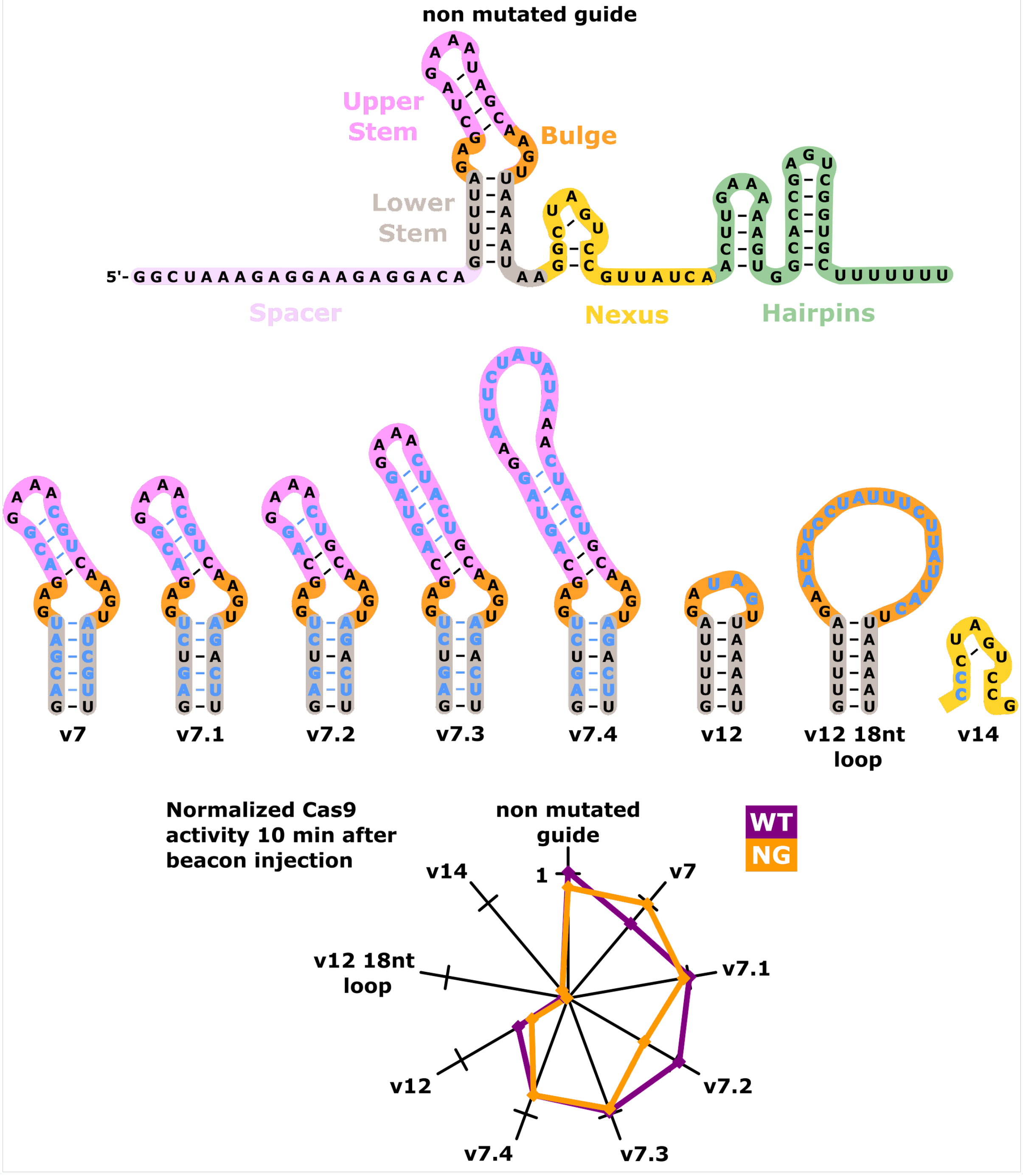

Supplementary Figure 3. Effect of mutations in the guide RNA's stem or nexus on the Cas9 activity. WT SpCas9 or SpCas9-NG was first incubated with full-length non-mutated guide RNA (top) or guide RNAs containing various mutations shown in blue in the schematic diagrams (the naming refers to the convention of Barrangou et al.). After stabilization of the fluorescence signal, the canonical beacon 2 (containing the PAM TGG) was injected and the signal amplitude was measured 10 min after injection. The signal amplitudes were normalized to the value obtained with WT SpCas9 in the presence of full-length non-mutated guide RNA and the canonical beacon 2. The radar chart shows the normalized activity (signal amplitude) of WT SpCas9 (purple) or SpCas9-NG (orange) in the presence of each guide RNA.

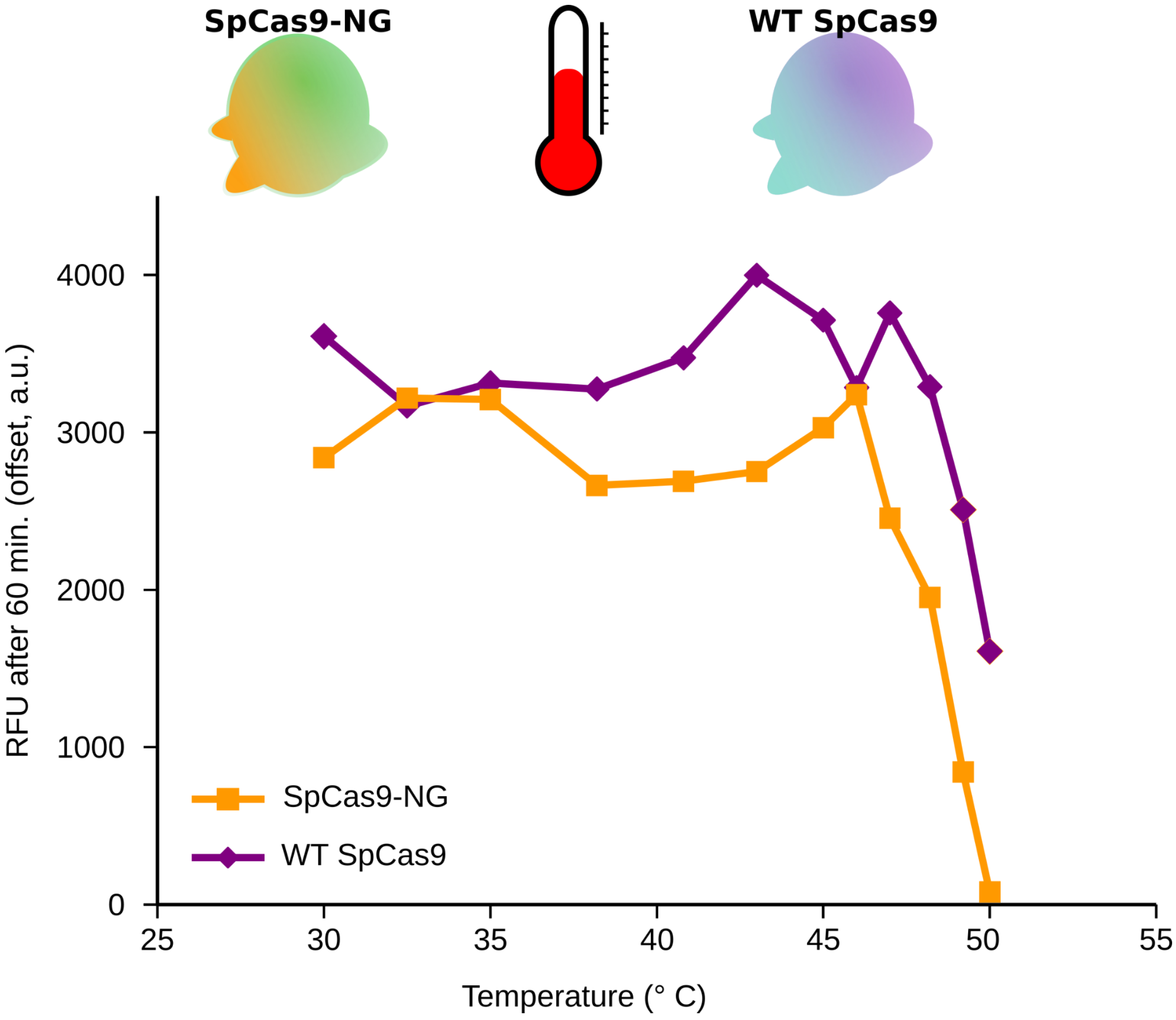

Supplementary Figure 4. Dependence of activity of Cas9 on temperature. The beacon 2 (40 nM, TGG PAM) was injected in a mixture of 200 nM of WT SpCas9 (purple) or SpCas9-NG (orange) pre-incubated with 400 nM of guide RNA for 30 minutes. The signal amplitudes are collected for each temperature 60 min after beacon injection.

| BEACONS | | | |
|---|---|---|---|
| NAME | SEQUENCE (5' -> 3') | MODIFICATIONS | |
| | | at 5' end | at 3' end |
| beacon 3 bottom CCA | ATTACGAATTCACCACTTTCCATTGGTCTCCTTCCTAT | | DY-681 |
| beacon 3 top TGG | ATAGGAAGGAGACCAATGGAAAGTGGTGAATTCGTAAT | BMN-Q650 | |
| beacon 3 bottom TCA | ATTACGAATTCATCACTTTCCATTGGTCTCCTTCCTAT | | DY-681 |
| beacon 3 top TGA | ATAGGAAGGAGACCAATGGAAAGTGATGAATTCGTAAT | BMN-Q650 | |
| beacon 3 bottom ACA | ATTACGAATTCAACACTTTCCATTGGTCTCCTTCCTAT | | DY-681 |
| beacon 3 top TGT | ATAGGAAGGAGACCAATGGAAAGTGTTGAATTCGTAAT | BMN-Q620 | |
| beacon 3 bottom GCA | ATTACGAATTCAGCACTTTCCATTGGTCTCCTTCCTAT | | DY-681 |
| beacon 3 top TGC | ATAGGAAGGAGACCAATGGAAAGTGCTGAATTCGTAAT | BMN-Q620 | |
| beacon 2 bottom CCA | ATTACGAATTCACCATGTCCTCTTCCTCTTTAGCCTAT | | Atto 647N |
| beacon 2 top TGG | ATAGGCTAAAGAGGAAGAGGACATGGTGAATTCGTAAT | BBQ-650 | |
| beacon 2 bottom TCA | ATTACGAATTCATCATGTCCTCTTCCTCTTTAGCCTAT | | DY-681 |
| beacon 2 top TGA | ATAGGCTAAAGAGGAAGAGGACATGATGAATTCGTAAT | BBQ-650 | |
| beacon 2 bottom ACA | ATTACGAATTCAACATGTCCTCTTCCTCTTTAGCCTAT | | Atto 590 |
| beacon 2 top TGT | ATAGGCTAAAGAGGAAGAGGACATGTTGAATTCGTAAT | BMN-Q620 | |
| beacon 2 bottom GCA | ATTACGAATTCAGCATGTCCTCTTCCTCTTTAGCCTAT | | Atto 532 |
| beacon 2 top TGC | ATAGGCTAAAGAGGAAGAGGACATGCTGAATTCGTAAT | BMN-Q535 | |

PAM sequences are underlined; sequences bound by the spacer (or their complementary sequence) are in purple

| GUIDE RNA TEMPLATES FOR BEACON 3 | |
|---|---|
| NAME | SEQUENCE (5' -> 3') |
| Template for reference guide | |
| beacon 3 sgRNA-103 | CACTACACTACTAACACACCACCAAATAATACGACTCACTATAGGAAGGAGACCAATGGAAAGGTTTTAGAGCTAGAAATAGCAAGTTAAAATAAGGCTAGTCCGTTATCAACTTGAAAAAGTGGCACCGAGTCGGTGCTTTTTTT |
| Templates for short guides | |
| beacon 3 sgRNA-20 | CGATGGACACAGACGCACGAACACGGCAACACGGCACATGCTACAGTTGCCTTACTTTGATTCACTACACTACTAACACACCACCAAATAATACGACTCACTATAGGAAGGAGACCAATGGAAAG |
| beacon 3 sgRNA-40 | ACACGGCAACACGGCACATGCTACAGTTGCCTTACTTTGATTCACTACACTACTAACACACCACCAAATAATACGACTCACTATAG |

| | |
|---|---|
| | GAAGGAGACCAATGGAAAGGTTTTAGAGCTAGAAATAGC |
| beacon 3 sgRNA-50 | ACGGCACATGCTACAGTTGCCTTACTTTGATTCACTACACTACTAACACACCACCAAATAATACGACTCACTATAGGAAGGAGACCAATGGAAAGGTTTTAGAGCTAGAAATAGCAAGTTAAAAT |
| beacon 3 sgRNA-62 | ACATGCTACAGTTGCCTTACTTTGATTCACTACACTACTAACACACCACCAAATAATACGACTCACTATAGGAAGGAGACCAATGGAAAGGTTTTAGAGCTAGAAATAGCAAGTTAAAATAAGGCTAGTCCG |
| beacon 3 sgRNA-63 | AGTTGCCTTACTCTTGATTCACTACACTACTAACACACCACCAAATAATACGACTCACTATAGGAAGGAGACCAATGGAAAGGTTTTAGAGCTAGAAATAGCAAGTTAAAATAAGGCTAGTCCGT |
| beacon 3 sgRNA-64 | AGTTGCCTTACTTTGATTCACTACACTACTAACACACCACCAAATAATACGACTCACTATAGGAAGGAGACCAATGGAAAGGTTTTAGAGCTAGAAATAGCAAGTTAAAATAAGGCTAGTCCGTT |
| beacon 3 sgRNA-65 | AGTTGCCTTACTTTGATTCACTACACTACTAACACACCACCAAATAATACGACTCACTATAGGAAGGAGACCAATGGAAAGGTTTTAGAGCTAGAAATAGCAAGTTAAAATAAGGCTAGTCCGTTA |
| beacon 3 sgRNA-66 | AGTTGCCTTACTTTGATTCACTACACTACTAACACACCACCAAATAATACGACTCACTATAGGAAGGAGACCAATGGAAAGGTTTTAGAGCTAGAAATAGCAAGTTAAAATAAGGCTAGTCCGTTAT |
| beacon 3 sgRNA-71 | AGTTGCCTTACTTTGATTCACTACACTACTAACACACCACCAAATAATACGACTCACTATAGGAAGGAGACCAATGGAAAGGTTTTAGAGCTAGAAATAGCAAGTTAAAATAAGGCTAGTCCGTTATCAACT |
| beacon 3 sgRNA-76 | TTGATTCACTACACTACTAACACACCACCAAATAATACGACTCACTATAGGAAGGAGACCAATGGAAAGGTTTTAGAGCTAGAAATAGCAAGTTAAAATAAGGCTAGTCCGTTATCAACTTGAAA |
| beacon 3 sgRNA-79 | TTGATTCACTACACTACTAACACACCACCAAATAATACGACTCACTATAGGAAGGAGACCAATGGAAAGGTTTTAGAGCTAGAAATAGCAAGTTAAAATAAGGCTAGTCCGTTATCAACTTGAAAAAG |
| beacon 3 sgRNA-80 | AGTTGCCTTACTTTGATTCACTACACTACTAACACACCACCAAATAATACGACTCACTATAGGAAGGAGACCAATGGAAAGGTTTTAGAGCTAGAAATAGCAAGTTAAAATAAGGCTAGTCCGTTATCAACTTGAAAAAGT |
| beacon 3 sgRNA-81 | AGTTGCCTTACTTTGATTCACTACACTACTAACACACCACCAAATAATACGACTCACTATAGGAAGGAGACCAATGGAAAGGTTTTAGAGCTAGAAATAGCAAGTTAAAATAAGGCTAGTCCGTTATCAACTTGAAAAAGTG |
| beacon 3 sgRNA-82 | CACTACACTACTAACACACCACCAAATAATACGACTCACTATAGGAAGGAGACCAATGGAAAGGTTTTAGAGCTAGAAATAGCAAGTTAAAATAAGGCTAGTCCGTTATCAACTTGAAAAAGTGG |
| beacon 3 sgRNA-90 | CACTACACTACTAACACACCACCAAATAATACGACTCACTATAGGAAGGAGACCAATGGAAAGGTTTTAGAGCTAGAAATAGCAAGTTAAAATAAGGCTAGTCCGTTATCAACTTGAAAAAGTGGCACCGAGT |
| beacon 3 sgRNA-100 | CACTACACTACTAACACACCACCAAATAATACGACTCACTATAGGAAGGAGACCAATGGAAAGGTTTTAGAGCTAGAAATAGCAAGTTAAAATAAGGCTAGTCCGTTATCAACTTGAAAAAGTGGCACCGAGTCGGTTTTTTT |
| Templates for mutated guides | |
| beacon 3 sgRNA-103 v3 | CACTACACTACTAACACACCACCAAATAATACGACTCACTATAGGAAGGAGACCAATGGAAAGGTCTCAGAGCTAGAAATAGCAAGTTGAGATAAGGCTAGTCCGTTATCAACTTGAAAAAGTGGCACC |

| | |
|---|---|
| | GAGTCGGTGCTTTTTTT |
| beacon 3 sgRNA-103 v4 | CACTACACTACTAACACACCACCAAATAATACGACTCACTATA GGAAGGAGACCAATGGAAAGGGGTTAGAGCTAGAAATAGCAAG TTAAAATAAGGCTAGTCCGTTATCAACTTGAAAAAGTGGCACC GAGTCGGTGCTTTTTTT |
| beacon 3 sgRNA-103 v12 | CACTACACTACTAACACACCACCAAATAATACGACTCACTATA GGAAGGAGACCAATGGAAAGGTTTTAGATACTTAAAATAAGGC TAGTCCGTTATCAACTTGAAAAAGTGGCACCGAGTCGGTGCTT TTTTT |
| beacon 3 sgRNA-103 v12 18nt loop | CACTACACTACTAACACACCACCAAATAATACGACTCACTATA GGAAGGAGACCAATGGAAAGGTTTTAGAATATCCTATTTCTTA TTACTTAAAATAAGGCTAGTCCGTTATCAACTTGAAAAAGTGG CACCGAGTCGGTGCTTTTTTT |
| beacon 3 sgRNA-103 v14 | CACTACACTACTAACACACCACCAAATAATACGACTCACTATA GGAAGGAGACCAATGGAAAGGTTTTAGAGCTAGAAATAGCAAG TTAAAATAACCCTAGTCCGTTATCAACTTGAAAAAGTGGCACC GAGTCGGTGCTTTTTTT |
| beacon 3 sgRNA-103 v15 ATTACT | CACTACACTACTAACACACCACCAAATAATACGACTCACTATA GGAAGGAGACCAATGGAAAGGTTTTAGAGCTAGAAATAGCAAG TTAAAATAAGACTAGTTCGATTACTACTTGAAAAAGTGGCACC GAGTCGGTGCTTTTTTT |
| Templates for guides with modified 3' hairpins | |
| beacon 3 sgRNA-103 no hp1 | CACTACACTACTAACACACCACCAAATAATACGACTCACTATA GGAAGGAGACCAATGGAAAGGTTTTAGAGCTAGAAATAGCAAG TTAAAATAAGGCTAGTCCGTTATCAA TGGCACCGAGTCGGTGCTTTTTTT |
| beacon 3 sgRNA-103 hp1-2p | CACTACACTACTAACACACCACCAAATAATACGACTCACTATA GGAAGGAGACCAATGGAAAGGTTTTAGAGCTAGAAATAGCAAG TTAAAATAAGGCTAGTCCGTTATCA TTGAAAAA GGCACCGAGTCGGTGCTTTTTTT |
| beacon 3 sgRNA-103 hp1-2d | CACTACACTACTAACACACCACCAAATAATACGACTCACTATA GGAAGGAGACCAATGGAAAGGTTTTAGAGCTAGAAATAGCAAG TTAAAATAAGGCTAGTCCGTTATCAAC GAAA GTGGCACCGAGTCGGTGCTTTTTTT |
| beacon 3 sgRNA-103 hp2-2d | CACTACACTACTAACACACCACCAAATAATACGACTCACTATA GGAAGGAGACCAATGGAAAGGTTTTAGAGCTAGAAATAGCAAG TTAAAATAAGGCTAGTCCGTTATCAACTTGAAAAAGTGGCAC AGT GTGCTTTTTTT |
| beacon 3 sgRNA-103 hp2-4d | CACTACACTACTAACACACCACCAAATAATACGACTCACTATA GGAAGGAGACCAATGGAAAGGTTTTAGAGCTAGAAATAGCAAG TTAAAATAAGGCTAGTCCGTTATCAACTTGAAAAAGTGGC AGT GCTTTTTTT |
| beacon 3 sgRNA-103 hp2-2p | CACTACACTACTAACACACCACCAAATAATACGACTCACTATA GGAAGGAGACCAATGGAAAGGTTTTAGAGCTAGAAATAGCAAG TTAAAATAAGGCTAGTCCGTTATCAACTTGAAAAAGTG ACCGAGTCGGT TTTTTTT |
| beacon 3 sgRNA-103 hp2-4p | CACTACACTACTAACACACCACCAAATAATACGACTCACTATA GGAAGGAGACCAATGGAAAGGTTTTAGAGCTAGAAATAGCAAG TTAAAATAAGGCTAGTCCGTTATCAACTTGAAAAAGTG CGAGTCG TTTTTTT |
| the spacer, stem, nexus and 3' hairpin sequences are respectively in purple, dark grey, orange, and green; mutations are in blue; deletions are empty spaces. | |

| GUIDE RNA TEMPLATES FOR BEACON 2 | |
|---|---|
| **NAME** | **SEQUENCE (5' -> 3')** |
| Template for reference guide | |
| beacon 2 sgRNA-103 | CACTACACTACTAACACACCACCAAATAATACGACTCACTATAGGCTAAAGAGGAAGAGGACAGTTTTAGAGCTAGAAATAGCAAGTTAAAATAAGGCTAGTCCGTTATCAACTTGAAAAAGTGGCACCGAGTCGGTGCTTTTTTT |
| Templates for short guides | |
| beacon 2 sgRNA-20 | CGATGGACACAGACGCACGAACACGGCAACACGGCACATGCTACAGTTGCCTTACTTTGATTCACTACACTACTAACACACCACCAAATAATACGACTCACTATAGGCTAAAGAGGAAGAGGACA |
| beacon 2 sgRNA-40 | ACACGGCAACACGGCACATGCTACAGTTGCCTTACTTTGATTCACTACACTACTAACACACCACCAAATAATACGACTCACTATAGGCTAAAGAGGAAGAGGACAGTTTTAGAGCTAGAAATAGC |
| beacon 2 sgRNA-50 | ACGGCACATGCTACAGTTGCCTTACTTTGATTCACTACACTACTAACACACCACCAAATAATACGACTCACTATAGGCTAAAGAGGAAGAGGACAGTTTTAGAGCTAGAAATAGCAAGTTAAAAT |
| beacon 2 sgRNA-56 | GAGCTAAGTGTACAATACGTACGCGAGTAGCACTTCCACTACACTACTAACACACCACCAAATAATACGACTCACTATAGGCTAAAGAGGAAGAGGACAGTTTTAGAGCTAGAAATAGCAAGTTAAAATAAGGCT |
| beacon 2 sgRNA-66 | AGTTGCCTTACTTTGATTCACTACACTACTAACACACCACCAAATAATACGACTCACTATAGGCTAAAGAGGAAGAGGACAGTTTTAGAGCTAGAAATAGCAAGTTAAAATAAGGCTAGTCCGTTAT |
| beacon 2 sgRNA-71 | AGTTGCCTTACTTTGATTCACTACACTACTAACACACCACCAAATAATACGACTCACTATAGGCTAAAGAGGAAGAGGACAGTTTTAGAGCTAGAAATAGCAAGTTAAAATAAGGCTAGTCCGTTATCAACT |
| beacon 2 sgRNA-76 | TTGATTCACTACACTACTAACACACCACCAAATAATACGACTCACTATAGGCTAAAGAGGAAGAGGACAGTTTTAGAGCTAGAAATAGCAAGTTAAAATAAGGCTAGTCCGTTATCAACTTGAAA |
| beacon 2 sgRNA-79 | TTGATTCACTACACTACTAACACACCACCAAATAATACGACTCACTATAGGCTAAAGAGGAAGAGGACAGTTTTAGAGCTAGAAATAGCAAGTTAAAATAAGGCTAGTCCGTTATCAACTTGAAAAAG |
| beacon 2 sgRNA-82 | CACTACACTACTAACACACCACCAAATAATACGACTCACTATAGGCTAAAGAGGAAGAGGACAGTTTTAGAGCTAGAAATAGCAAGTTAAAATAAGGCTAGTCCGTTATCAACTTGAAAAAGTGG |
| beacon 2 sgRNA-90 | CACTACACTACTAACACACCACCAAATAATACGACTCACTATAGGCTAAAGAGGAAGAGGACAGTTTTAGAGCTAGAAATAGCAAGTTAAAATAAGGCTAGTCCGTTATCAACTTGAAAAAGTGGCACCGAGT |
| Templates for mutated guides | |
| beacon 2 sgRNA-82 v3 | CACTACACTACTAACACACCACCAAATAATACGACTCACTATAGGCTAAAGAGGAAGAGGACAGTCTCAGAGCTAGAAATAGCAAGTTGAGATAAGGCTAGTCCGTTATCAACTTGAAAAAGTGG |
| beacon 2 sgRNA-82 v4 | CACTACACTACTAACACACCACCAAATAATACGACTCACTATAGGCTAAAGAGGAAGAGGACAGGGTTAGAGCTAGAAATAGCAAGTTAAAATAAGGCTAGTCCGTTATCAACTTGAAAAAGTGG |
| beacon 2 sgRNA-82 v7 | CACTACACTACTAACACACCACCAAATAATACGACTCACTATAGGCTAAAGAGGAAGAGGACAGACGATGAGACGGAAACGTCAAG |

| | |
|---|---|
| | TATCGTTAAGGCTAGTCCGTTATCAACTTGAAAAAGTGG |
| beacon 2<br>sgRNA-82 v15<br>ATTACT | CACTACACTACTAACACACCACCAAATAATACGACTCACTATA<br>GGCTAAAGAGGAAGAGGACAGTTTTAGAGCTAGAAATAGCAAG<br>TTAAAATAAGACTAGTTCGATTACTACTTGAAAAAGTGG |
| beacon 2<br>sgRNA-103 v7 | CACTACACTACTAACACACCACCAAATAATACGACTCACTATA<br>GGCTAAAGAGGAAGAGGACAGACGATGAGACGGAAACGTCAAG<br>TATCGTTAAGGCTAGTCCGTTATCAACTTGAAAAAGTGGCACC<br>GAGTCGGTGCTTTTTTT |
| beacon 2<br>sgRNA-103 v7.1 | CACTACACTACTAACACACCACCAAATAATACGACTCACTATA<br>GGCTAAAGAGGAAGAGGACAGAGTCTGAGACGGAAACGTCAAG<br>TAGACTTAAGGCTAGTCCGTTATCAACTTGAAAAAGTGGCACC<br>GAGTCGGTGCTTTTTTT |
| beacon 2<br>sgRNA-103 v7.2 | CACTACACTACTAACACACCACCAAATAATACGACTCACTATA<br>GGCTAAAGAGGAAGAGGACAGAGTCTGAGCAGGAAACTGCAAG<br>TAGACTTAAGGCTAGTCCGTTATCAACTTGAAAAAGTGGCACC<br>GAGTCGGTGCTTTTTTT |
| beacon 2<br>sgRNA-103 v7.3 | CACTACACTACTAACACACCACCAAATAATACGACTCACTATA<br>GGCTAAAGAGGAAGAGGACAGAGTCTGAGCAGTAGGAAACTAC<br>TGCAAGTAGACTTAAGGCTAGTCCGTTATCAACTTGAAAAAGT<br>GGCACCGAGTCGGTGCTTTTTTT |
| beacon 2<br>sgRNA-103 v7.4 | CACTACACTACTAACACACCACCAAATAATACGACTCACTATA<br>GGCTAAAGAGGAAGAGGACAGAGTCTGAGCAGTAGGAATTCTA<br>TATAAACTACTGCAAGTAGACTTAAGGCTAGTCCGTTATCAAC<br>TTGAAAAAGTGGCACCGAGTCGGTGCTTTTTTT |
| beacon 2<br>sgRNA-103 v12 | CACTACACTACTAACACACCACCAAATAATACGACTCACTATA<br>GGCTAAAGAGGAAGAGGACAGTTTTAGATACTTAAAATAAGGC<br>TAGTCCGTTATCAACTTGAAAAAGTGGCACCGAGTCGGTGCTT<br>TTTTT |
| beacon 2<br>sgRNA-103 v12<br>18nt loop | CACTACACTACTAACACACCACCAAATAATACGACTCACTATA<br>GGCTAAAGAGGAAGAGGACAGTTTTAGAATATCCTATTTCTTA<br>TTACTTAAAATAAGGCTAGTCCGTTATCAACTTGAAAAAGTGG<br>CACCGAGTCGGTGCTTTTTTT |
| beacon 2<br>sgRNA-103 v14 | CACTACACTACTAACACACCACCAAATAATACGACTCACTATA<br>GGCTAAAGAGGAAGAGGACAGTTTTAGAGCTAGAAATAGCAAG<br>TTAAAATAACCCTAGTCCGTTATCAACTTGAAAAAGTGGCACC<br>GAGTCGGTGCTTTTTTT |

the spacer, stem, nexus and 3' hairpin sequences are respectively in purple, dark grey, orange, and green; mutations are in blue.

Table 1: DNA sequences of fluorescence reporters and DNA templates for *in vitro* transcription
